# Population size determines the type of nucleotide variations in humans

**DOI:** 10.1101/130294

**Authors:** Sankar Subramanian

## Abstract

It is well known that the effective size of a population (*N*_*e*_) is one of the major determinants of the amount of genetic variation within the population. Here, we examined whether the types of genetic variations are dictated by the effective population size. Our results revealed that for low frequency variants, the ratio of AT→GC to GC→AT variants (*β*) was similar across populations, suggesting the similarity of the pattern of mutation in various populations. However, for high frequency variants, *β* showed a positive correlation with the effective population size of the populations. This suggests a much higher proportion of high frequency AT→GC variants in large populations (e.g. Africans) compared to those with small population sizes (e.g. Asians). These results imply that the substitution patterns vary significantly between populations. These findings could be explained by the effect of GC-biased gene conversion (gBGC), which favors the fixation of G/C over A/T variants in populations. In large population, gBGC causes high *β*. However, in small populations, genetic drift reduces the effect of gBGC resulting in reduced *β*. This was further confirmed by a positive relationship between *N*_*e*_ and *β* for homozygous variants. Our results highlight the huge variation in the types of homozygous and high frequency polymorphisms between world populations. We observed the same pattern for deleterious variants, implying that the homozygous polymorphisms associated with recessive genetic diseases will be more enriched with G or C in populations with large *N*_*e*_ (e.g. Africans) than in populations with small *N*_*e*_ (e.g. Europeans).

## Introduction

The out of Africa hypothesis predicts that the ancestors of the human populations around the world originated in Africa, migrated out of the continent and eventually colonized different parts of the world (Stringer 2003). During this process, the ancestral populations underwent a series of population bottlenecks along the migratory routes. Due to this founder effect, the ancestral population size is expected to decline with increasing distance from Africa. Previous empirical studies confirmed this prediction and showed that populations in Africa are the most genetically diverse and that the diversity declined with increasing geographic distance from Africa particularly along the colonization routes (Prugnolle, et al. 2005; Ramachandran, et al. 2005; Handley, et al. 2007; Li, et al. 2008; DeGiorgio, et al. 2009). These observations clearly suggest significant variation in the nucleotide diversity among global populations.

Nucleotide diversity (*π=*4*N*_*e*_*μ*) is a measure of genetic variation, which is determined by mutation rate (*μ*) and effective population size (*N*_*e*_). Since mutation rate is similar across human populations, the observed difference in the diversity of world populations is largely due to the variations in effective population sizes. Although a recent study suggested a higher rate of mutation in non-Africans, the magnitude of this effect was very small (∼5%) (Mallick, et al. 2016). Recent population genomic studies showed large variation in the number of polymorphisms observed between world populations (Genomes Project, et al. 2015). Populations in Africa have ∼5 million Single Nucleotide Variations (SNVs) whereas those in East Asia have ∼4.1 million, which is ∼20% less. Although the variation in the number of polymorphisms is well known, it is unclear if there are differences in the types of polymorphisms between world populations. Are the frequencies of different types of nucleotide changes (eg. A→G or T→C) similar across populations? This question arises from our understanding of the phenomenon of GC-biased gene conversion (gBGC). gBGC is a recombination-associated process that favors G/C over A/T nucleotides during the repair of mismatches that occur in heteroduplex DNA during meiosis (Duret, et al. 2002; Marais 2003; Duret and Galtier 2009; Galtier, et al. 2009; Glemin, et al. 2015). Although this process is not associated with natural selection, the efficiency of gBGC could also be reduced by genetic drift. Therefore, the effect of gBGC is expected to be much weaker in small populations than in large populations (Galtier, et al. 2009; Glemin, et al. 2015) and consequently, the frequencies of AT→GC polymorphisms are expected to differ among world populations. It is important to characterize the population specific patterns of genetic variants, as these patterns may have immense implications for human health.

For instance, previous studies have shown a positive correlation between the number of deleterious homozygous SNVs present in human populations and their distance from East Africa (Henn, et al. 2016). Furthermore, non-Africans were found to have much higher proportion of high frequency or homozygous deleterious variants than Africans (Do, et al. 2015; Subramanian 2016). These observations suggest that due to the effects of genetic drift, small populations (typically located away from Africa) have a higher fraction of high frequency (and homozygous) deleterious mutations than large populations. On the other hand, a gBGC mediated skews in the frequencies of deleterious AT→GC (relative to GC→AT) polymorphisms were also reported (Lachance and Tishkoff 2014; Xue, et al. 2016). However, it is unclear whether the extent of such skews is influenced by the effective population sizes of various global populations. Therefore, using data from the 1000 Genomes Project we investigated the pattern of nucleotide changes observed in the SNVs segregating in different allele frequencies (Genomes Project, et al. 2015). Furthermore, we also analyzed homozygous and heterozygous variant data for 126 distinct populations from around the world, obtained from the Simons Genome Diversity Project (Mallick, et al. 2016).

## Results

### Allele frequencies and types of nucleotide changes in human populations

To quantify the difference in the patterns of observed AT→GC and GC→AT changes we derived a measure *β*, based on the Waterston estimator (*θ*_*W*_) as described in the methods (Equation 1). The measure *β* is the ratio of AT→GC (*μ*_*AT→GC*_) and GC→AT (*μ*_*GC→AT*_) mutation rates, which captures the mutational equilibrium between AT and GC nucleotides. The ratio *β* is expected to be 1 if the observed AT→GC and GC→AT changes are due to the forward and reverse mutation rates alone. Any deviation from this ratio (*β*=1) suggests a bias in the substitution of one type over the other. To examine the variation in the patterns of nucleotide changes we obtained the 1000 Genomes phase II data for 26 distinct populations of the world. Since most of the genomes from Latin America were admixed with Europeans/Africans, we separated 20 Peruvian genomes that had <0.5% admixture. We used these to represent an un-admixed Native American population and hence the total number of populations analyzed was 27. The SNVs of these populations were grouped into eleven categories based on their Derived Allele Frequencies (DAF). The ratio *β* was estimated for each category of SNVs belonging to each population. We estimated the effective size (*N*_*e*_) of each population based on the mutation rate (*μ*) and nucleotide diversity (*π*) (see methods).

Figure 1 shows the relationship between *N*_*e*_ and *β* for SNVs belonging to two extreme allele frequencies. For DAF < 0.025, the estimates of *β* were almost equal to 1 for all populations. In contrast, we observed a significant positive correlation (*P* < 10^-6^) for SNVs with very high DAF (> 0.9). Another major pattern was that the *β* estimates were similar among populations belonging to the same geographical locations and very different among populations from distinct locations. However, this was not true for admixed Americans (black dots). We then examined this relationship for SNVs with different derived allele frequencies. We did not find any significant relationship between *N*_*e*_ and *β* for low frequency SNVs until the DAF was ≤ 0.2 (*P* > 0.10) (Figure 2A). However, significant positive correlations (at least *P* < 10^-2^) were observed between *N*_*e*_ and *β* for SNVs with DAF > 0.2. The magnitude of the correlation increased with the increase in DAF, which is evident from the rise in the slopes of the regression lines. This increase is clearer in Figure 2B, which shows the positive relationship between DAF and the slopes of the regression lines shown in Figure 2A. The slope observed for SNVs with DAF > 0.9 was 6 ×10^-5^, which is almost 15 times higher than for SNVs with DAF = 0.2-0.3 (4 ×10^-6^). For high frequency SNVs (DAF > 0.9), the difference between the mean *β* estimated for African genomes (1.72) was 79% higher than that observed for Peruvian genomes (1.35).

**Figure 1.**
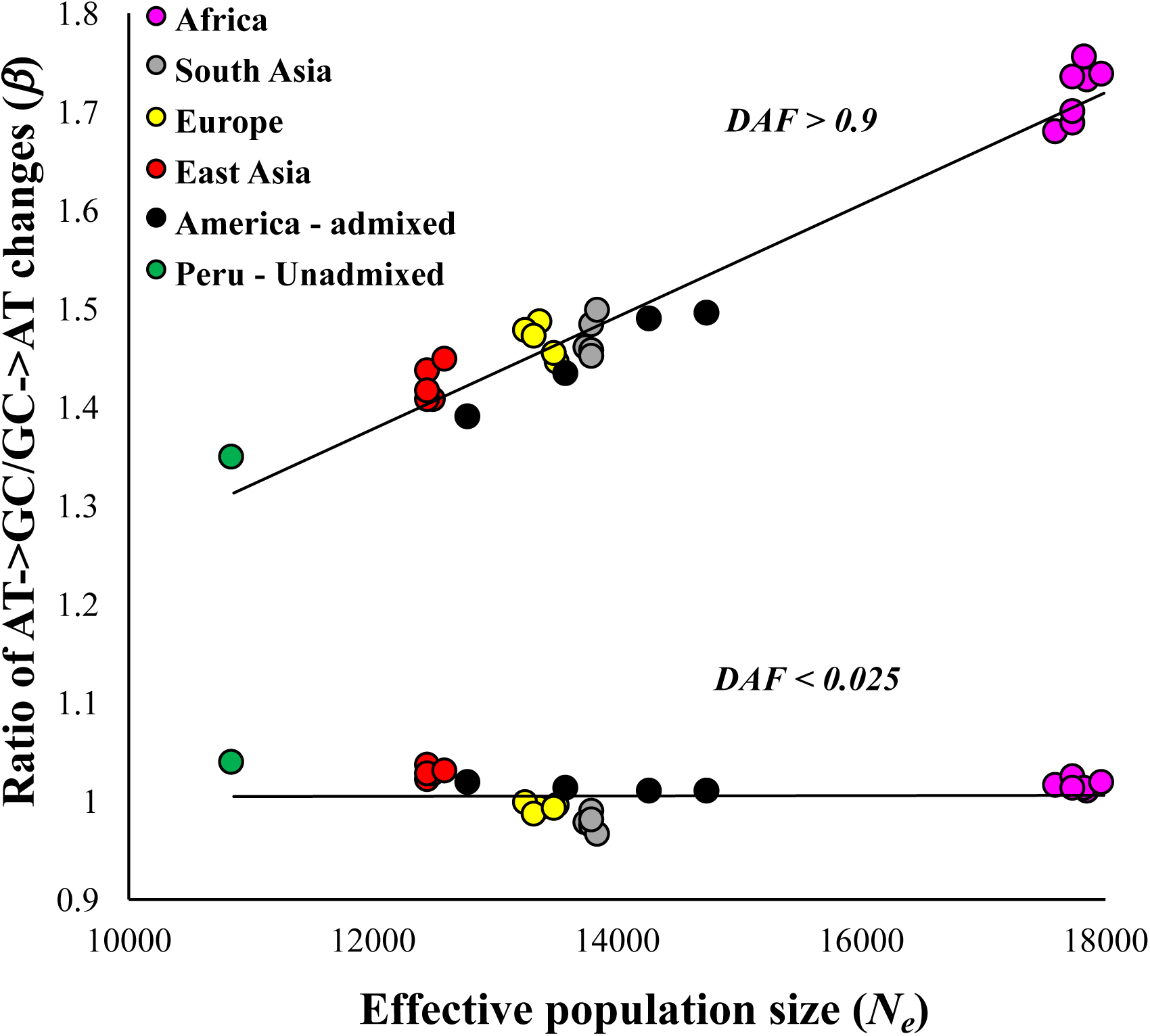
Relationship between the effective population size (*N*_*e*_) and the ratio of AT→GC to GC→AT (*β*) changes. The genotype data was obtained for 27 populations of the world. The colors indicate different geographical locations of the populations. The relationship was not significant for SNVs with DAF <0.025 (*P* = 0.11, using the Spearman rank correlation) but highly significant for those with DAF > 0.9 (*P* < 10^-6^).

**Figure 2.**
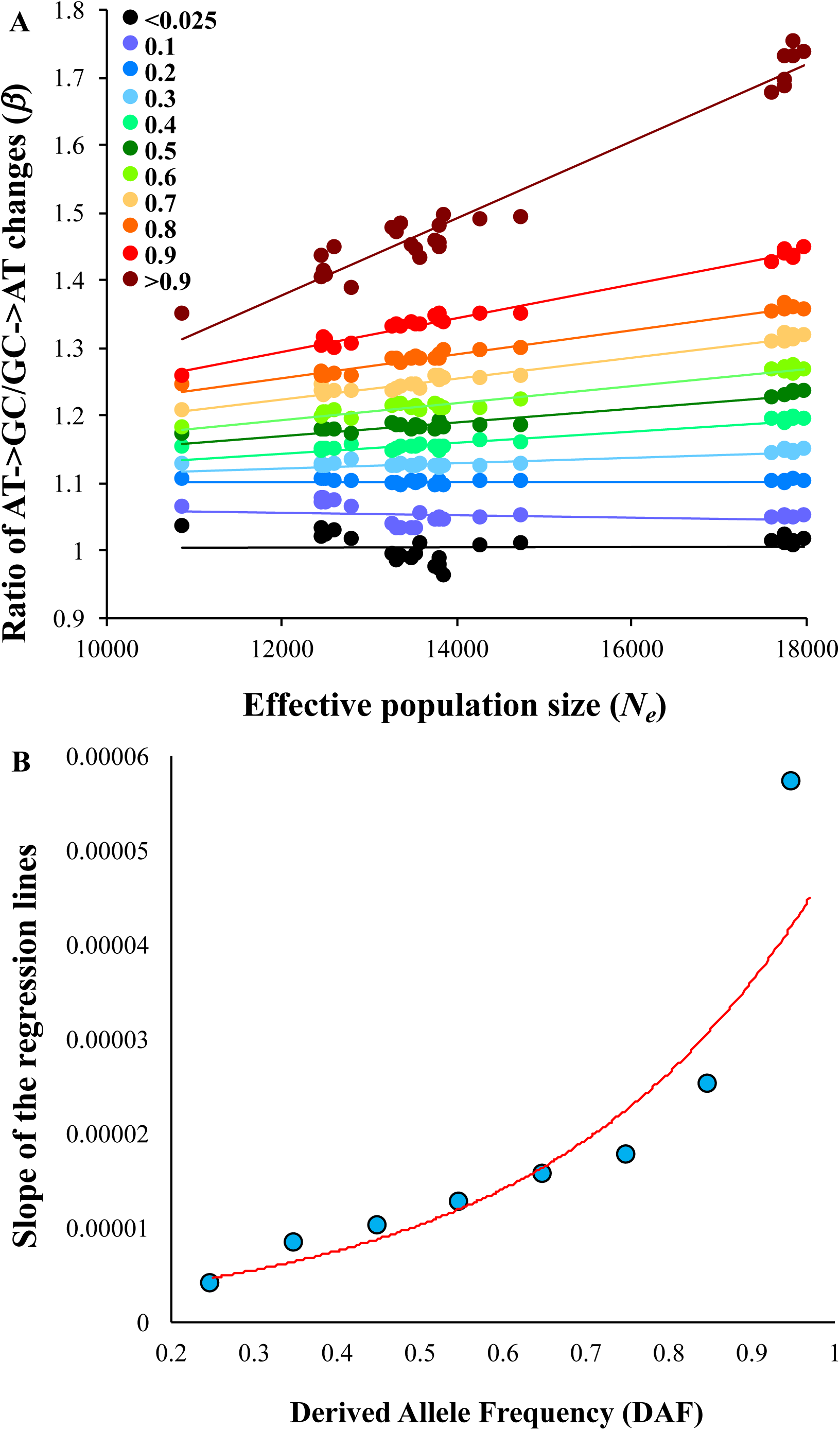
**(A)** The correlation between *N*_*e*_ and *β* observed for SNVs belonging to eleven allele frequency categories with DAF: <0.025, 0.025-0.1, 0.1-0.2, 0.2-0.3, 0.3-0.4, 0.4-0.5, 0.5-0.6, 0.6-0.7, 0.7-0.8, 0.8-0.9 and > 0.9. Only the upper values of the range are shown in X-axis. The correlations observed for SNVs with DAF < 0.2 were not statistically significant (P > 0.10) and remaining relationships were highly significant (at least P < 10^-2^). **(B)** Scatter plot showing the magnitude of the relationship shown in figure 2A for difference allele frequency categories. X-axis shows the mid-values of the derived allele frequency categories and the slopes of the regression lines of figure 2A are shown on the Y-axis. The slopes of the lines for SNVs with DAF ≤ 0.2 are not included as those relationships were not statistically significant.

We also plotted *β* against DAF to show how this estimate changes with increasing DAF. For this purpose, we selected five representative populations with significantly different *N*_*e*_. As shown in Figure 3, positive relationships between DAF and *β* against were observed for all populations and *β* increased with increasing DAF. However, based on the slopes of the regression lines (see Figure 2 legend-inset) the rate of increase correlates with *N*_*e*_. The slope was the highest for Africans (0.57), intermediate for Eurasians (0.44-0.32) and the lowest for Peruvians (0.27).

**Figure 3.**
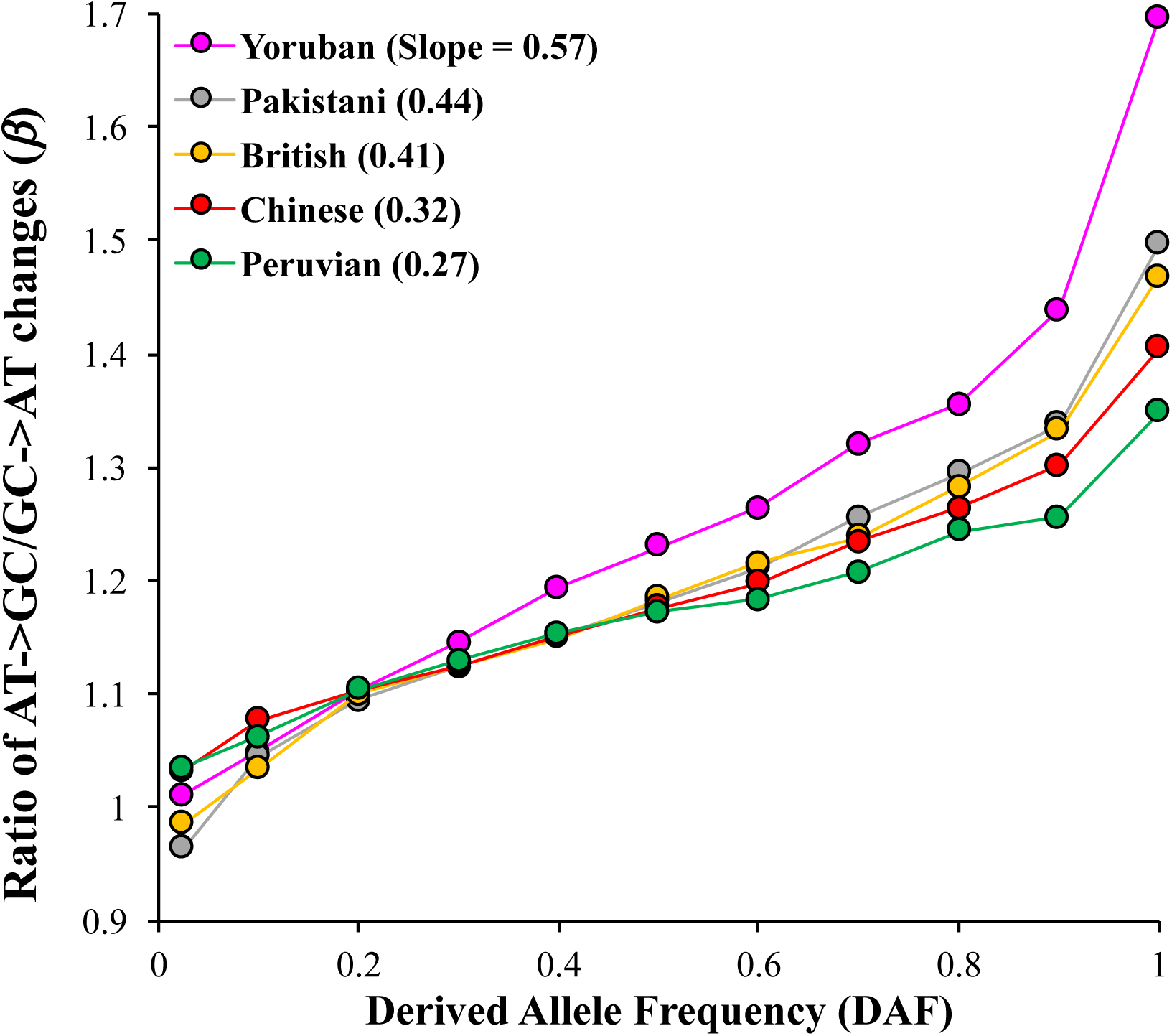
The magnitude of *β* with respect to the derived allele frequencies. The correlation between DAF and *β* were highly significant (at least P < 0.0017). Although all populations show an increasing trend, the rate of increase varies between them, which are manifested in the slopes of the regression lines (shown in the figure legend).

### Patterns of homozygous and heterozygous variations

To further examine the patterns of substitutions in a much larger number of populations we obtained the genotype data of the Simons Genome Diversity Project for genomes representing 126 distinct ethnic populations. We examined the patterns of nucleotide changes for homozygous and heterozygous SNVs. For each genome, we estimated *β* for homozygous and heterozygous SNVs. The nucleotide diversity was estimated by comparing the pairs of chromosomes of each genome and *N*_*e*_ was calculated using the mutation rate obtained from previous studies (see methods). We then plotted *β* against *N*_*e*_. For homozygous SNVs, our regression analysis produced a significant positive correlation (*P* < 10^-6^) between the two variables (Figure 4A). Although the effective population sizes vary widely between world populations, they are roughly similar for the populations in a geographical location. Interestingly, the *β* values for homozygous SNVs are also varied considerably but were similar for populations within a geographical region. This is clear from the mean estimates shown in the inset of Figure 4A. On the other hand, *N*_*e*_ for heterozygous SNVs did not show any significant relationship (*P* = 0.36) with *β* (Figure 4B).

**Figure 4.**
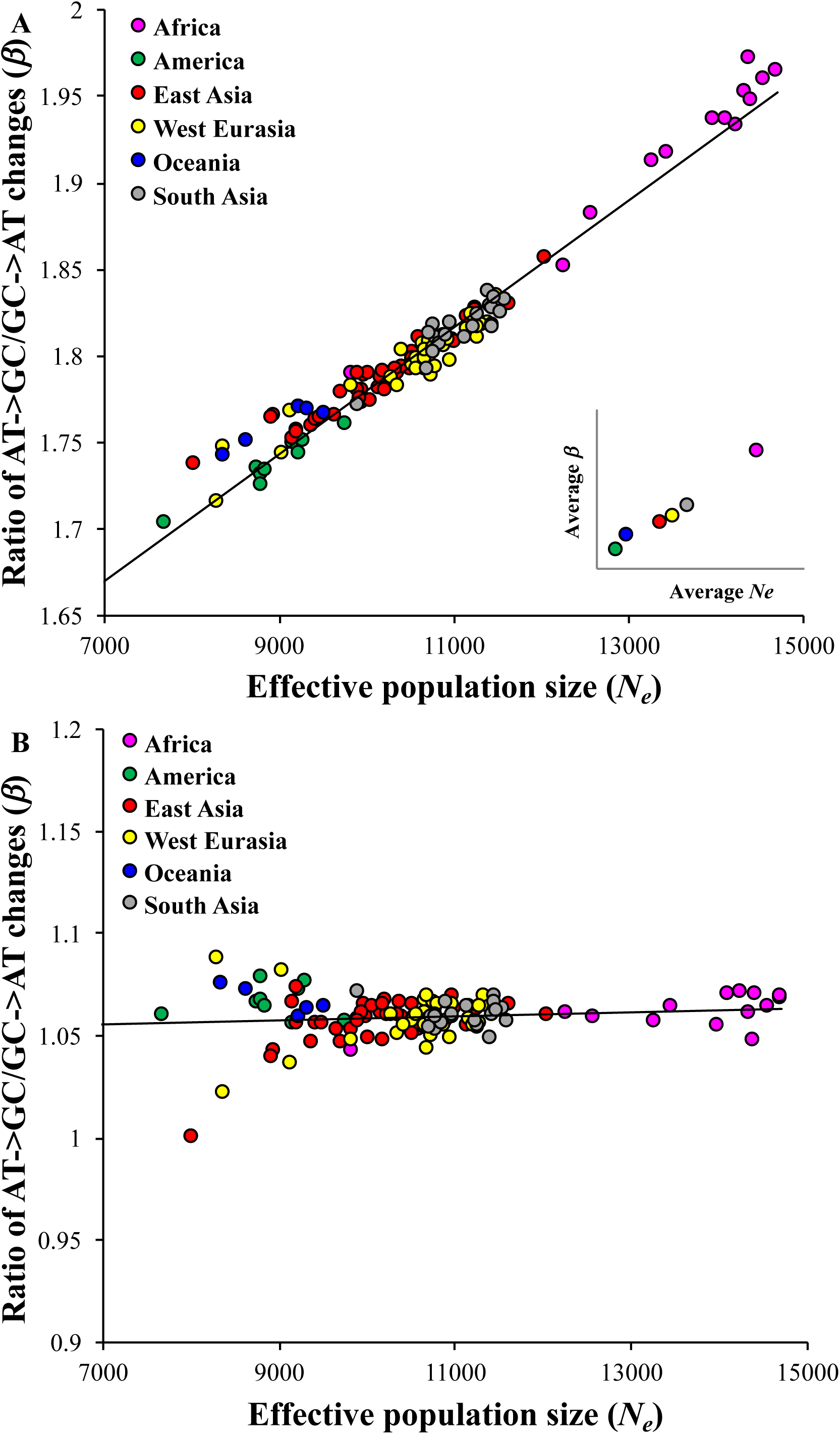
The relationship between effective population size (*N*_*e*_) and the normalized ratio AT→GC to GC→AT (*β*) estimated for homozygous **(A)** and heterozygous **(B)** SNVs present in individual genomes belonging to 126 distinct populations of the world. The data was obtained from the Simons Genome Variation Project. The correlation for homozygous SNVs was highly significant (*P* < 10^-6^) but not statistically significant for heterozygous SNVs (*P* = 0.36). The colors represent populations from distinct geographical locations. The relationship shown in the inset was based on the estimates of *N*_*e*_ and *β* averaged for the populations belonging to a specific geographical location. The relationship shown in the inset was also significant (P < 0.026).

### Proportion of deleterious AT→GC and GC→AT changes in global populations

To understand the implications of the observed patterns on human health, we examined the patterns of nucleotide changes for deleterious SNVs. To determine the deleteriousness of the SNVs we used the robust method, Combined Annotation-Dependent Depletion (CADD). This method integrates over 60 diverse annotations (to measure the extent of deleteriousness of a variant) into a single measure (*C* score) (Kircher, et al. 2014). We designated SNVs with a *C* score of >15 as deleterious. To distinguish the frequency of different of types of nucleotide changes we estimated the proportion of deleterious AT→GC (*P*_*AT*_) and GC→AT (*P*_*GC*_) changes using equations 2 and 3 (methods). These estimates were plotted against *N*_*e*_. For deleterious SNVs with DAF > 0.9 we found highly significant relationships between *N*_*e*_ and *P*_*GC*_ (*P* < 10^-6^) and between *N*_*e*_ and *P*_*AT*_ (*P* < 10^-6^) (Figure 5A). Importantly for large populations (Africans) the proportion of deleterious AT→GC mutations was 54% higher than the proportion of GC→AT mutations. For small populations (Peruvians) this difference was only 26%. We performed a similar analysis using the homozygous SNVs from 126 world populations and obtained similar results (Figure 5B). The difference between *P*_*GC*_ and *P*_*AT*_ was highest for Africans (51%) and lowest (22%) for Native Americans.

**Figure 5.**
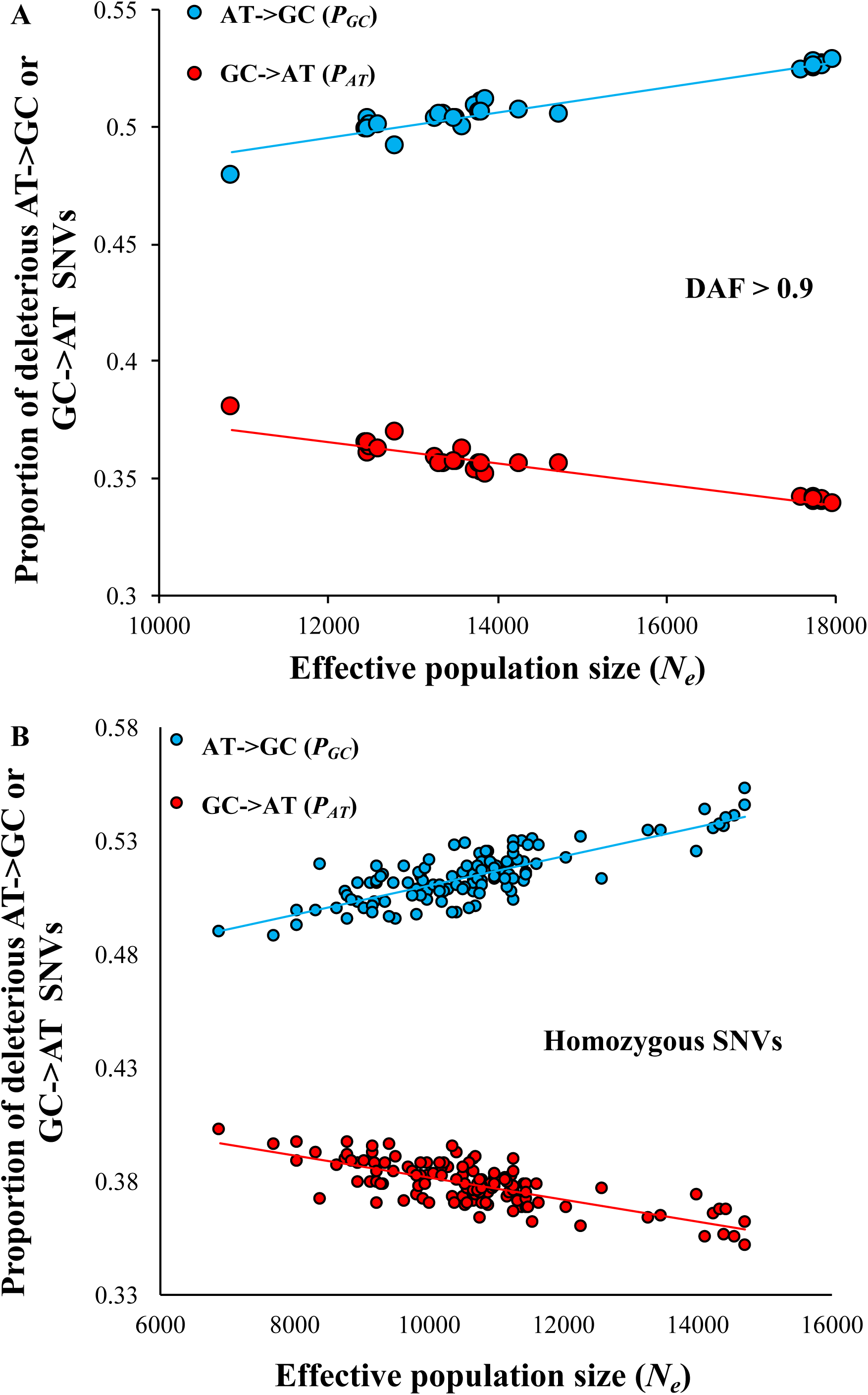
Variations in the proportions of deleterious AT→GC and GC→AT SNVs among world populations. These proportions correlate significantly with the effective population sizes of the populations. The trends were positive for AT→GC SNVs and negative for GC→AT SNVs. The relationships were based on **(A)** high frequency SNVs with DAF > 0.9 and **(A)** homozygous SNVs. All relationships were highly significant (at least, *P* < 10^-6^).

## Discussions

In this study, we showed that the types of nucleotide changes in world populations are shaped by their effective population sizes. Our results revealed a much higher proportion of AT→GC variations in populations with large effective sizes (eg. Africans) compared to those with small sizes (eg. Native Americans). These observations could be explained based on the well-known recombination associated GC-biased gene conversion (gBGC) (Duret, et al. 2002; Marais 2003; Duret and Galtier 2009; Galtier, et al. 2009; Glemin, et al. 2015). The two strands of DNA are connected by double hydrogen bonds between A and T bases (weak) as well as triple hydrogen bonds between G and C bases (strong). It has been shown that gBGC favors the changes involving AT→GC (or weak → strong) compared to GC→AT (or strong → weak) during the process of fixation. We developed a measure *β* to capture this bias in substitutions. For rare SNVs, *β* estimates were close to 1 (*β*≈1) for all populations (Figure 1). This suggests that the rate of AT→GC mutations per (A or T) base is equal to GC→AT mutations per (G or C) base. Therefore, the observed number of rare SNVs reflect the mutation patterns alone.

In contrast, for high frequency variations *β* estimates were significantly higher than 1 (*β* > 1). Importantly *β* increased with the increase in DAF as shown clearly in Figure 3. These results suggest a preferential fixation of AT→GC over GC→AT mutations over time. However, based on the slopes of the regression lines the results also suggest that the rate of preferential of fixation of AT→GC mutations in small populations is low compared to that in large populations because of genetic drift that reduces the efficiency of gBGC. This result is supported by a previous study using human population genetic data that suggested that on an average gBGC is stronger in African than non-African populations (Glemin, et al. 2015).

To further support our claim, we examined the changes between A↔T (weak bond) and between G↔C (strong bond) nucleotides using high frequency SNVs (DAF > 0.9). Our results showed that *β* estimated for A→T/T→A or C→G/G→C were close to 1 for all populations and there was no significant relationship with *N*_*e*_ (Supplementary Figure S1). We also examined this using homozygous and heterozygous SNVs from the 126 populations and observed no significant relationship between *β* and *N*_*e*_ (Supplementary Figure S2A and S2B). Since the changes are between the same types of nucleotides (with respect to weak or strong hydrogen bonds) there was no effect of gBGC on the fixation of one type of nucleotide over other. This provides further evidence that the results of our study are not due to methodological artifacts.

Since gBGC does not affect the changes between A↔T (weak bond) and between G↔C (strong bond) previous studies have used the rate of these changes as a normalizing factor to assess the magnitude of gBGC on GC↔AT changes (Lachance and Tishkoff 2014; Glemin, et.al 2015; Xue and Chen 2016). Following this, we normalized AT→GC with A↔T changes and GC→AT with G↔C changes respectively and developed a normalized ratio, *β’* (equation 2 - methods). However, the relationship between *N*_*e*_ and *β’* was also highly significant and comparable to previous results obtained for high frequency (Supplementary Figure S3A) and homozygous SNVs (Supplementary Figure S3B). This further supports our results as the normalization eliminates any variation in the mutation rates between populations.

The results based on homozygous and heterozygous SNVs shown in Figure 4 further support to the results based on DAF presented in Figures 1 and 2. For instance, Figures 1 and 2A showed there was no significant relationship between *β* and *N*_*e*_ for low frequency SNVs. Since low frequency SNVs predominantly exist as heterozygous SNVs in a genome, results based on the former are expected to be similar to those based on the latter (Figures 1 and Figure 4B). Similarly, as high frequency SNVs are more likely to be present as homozygous SNVs in genomes, the results for these two types of SNVs are alike. This is evident from the results shown in Figures 1 and 4A.

Since gBGC is mediated by recombination it effects were found to be strong for highly recombining regions. To examine this, we obtained SNVs from regions with low (<2 cM), medium (2-20 cM) and high (>20 cM) rates of recombination. The results showed a much higher *β* for the variants in high recombination regions (Supplementary Figure S4). However, the magnitude of the relationship between *β* and *N*_*e*_ was similar for all three regions; the slopes of the regression lines were 0.000039 (low), 0.000042 (medium) and 0.000041 (high). This suggests that the influence of population size on gBGC is relatively similar across chromosomal regions with varying degrees of recombination.

Previous studies have shown a significant negative correlation between heterozygosity and the distance (of the location of the populations) from Africa. This correlation is expected based on the prediction that during migration out of Africa human populations underwent a series of population bottlenecks or founder effects along the migratory route (Prugnolle, et al. 2005; Ramachandran, et al. 2005; Handley, et al. 2007; DeGiorgio, et al. 2009). This is because only a subset of people migrated from the original location to new sites and hence the size of the populations reduced with distance from Africa. From previous studies, we obtained the geographic distance of 41 non-African populations from Eastern Africa (Addis Ababa) (Li, et al. 2008; DeGiorgio, et al. 2009) and we plotted the estimates of *β* against them. We obtained a highly significant negative correlation for homozygous variants (*P* < 10^-6^) (Figure 6A) but not for heterozygous variants (*P* = 0.2) (Figure 6B). This is very similar to the results shown in Figure 4. In this analysis, the geographic distance from Africa was used as a proxy for *N*_*e*_. Hence this result independently confirms our findings and also justifies the method used in this study to estimate *N*_*e*_ from nucleotide diversity.

**Figure 6.**
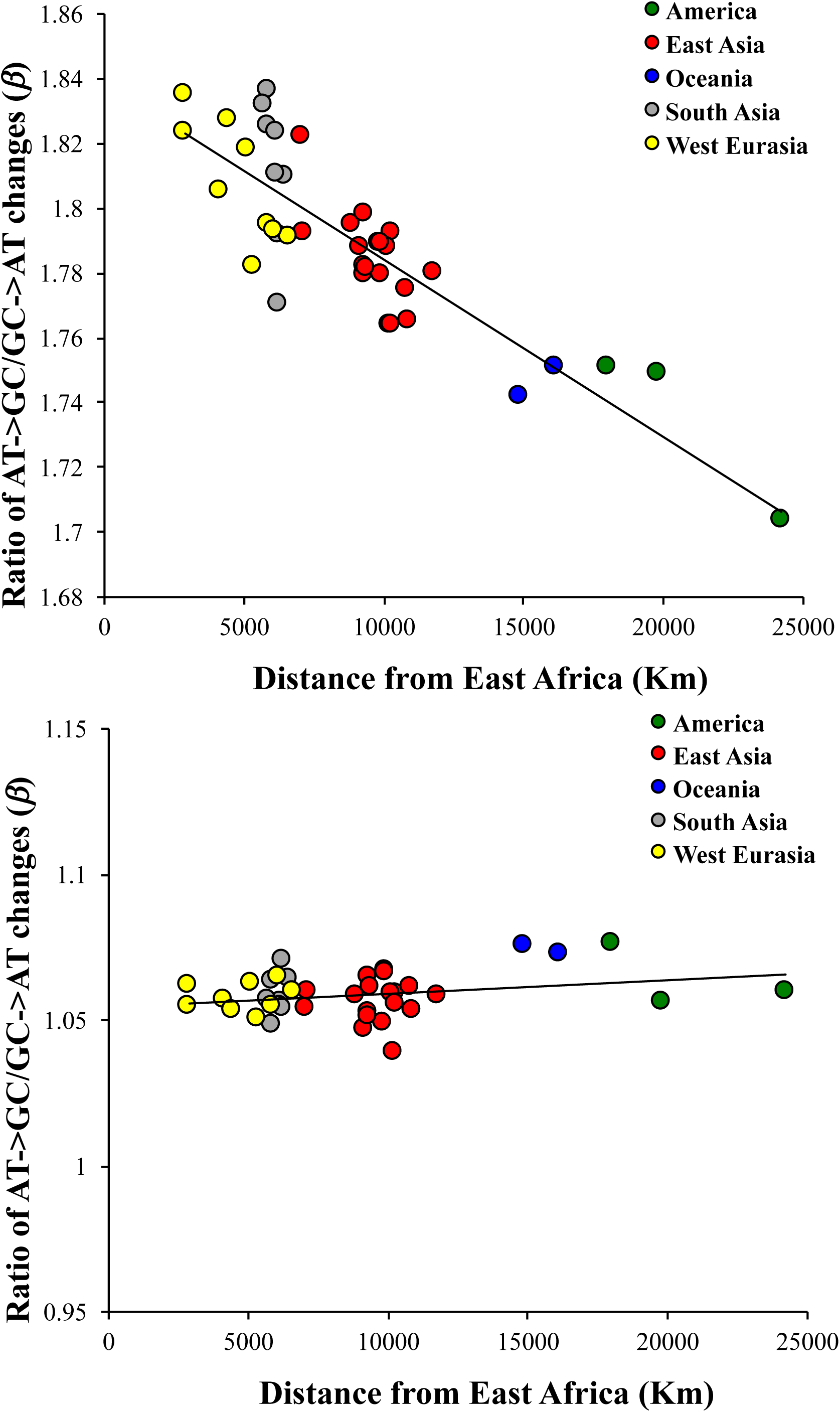
Correlation between the geographical distance of non-African populations from East Africa (Addis Ababa) and the normalized ratio AT→GC/GC→AT (*β*) using **(A)** Homozygous **(B)** Heterozygous SNVs. The relationship was highly significant for homozygous SNVs (*P* < 10^-6^) and not for heterozygous SNVs (*P* = 0.2). The distances between the locations of the non-African populations and Addis Ababa were obtained from previous studies (Li, et al. 2008; DeGiorgio, et al. 2009).

We estimated *N*_*e*_ under the assumption that mutation rate (*μ*) and the rate of accumulation of mutations are both similar between populations. However, a recent study suggested a slightly (∼5%) higher rate of mutation accumulation in non-Africans compared to Africans (Mallick, et al. 2016). To accommodate the elevated diversity, we subtracted 5% of the observed divergence for non-Africans while estimating *N*_*e*_ and re-analyzed the data. This produced almost identical patterns and similar strengths of correlation to that reported in Figure 4 (Supplementary Figure S5).

We have shown that the types of SNVs observed in different human populations are very likely to be modulated by their effective population sizes. Since this pattern was universal for genome-wide variations we showed that deleterious SNVs also follow this pattern. Our results showed that populations with large effective sizes (e.g. Africans) displayed the greatest difference between the proportions of high frequency deleterious AT→GC and GC→AT SNVs. This difference was much lower in populations with small effective sizes (e.g. Native Americans). This has significant implications in human health as it implies that high frequency diseases-associated mutations in Africans will be more enriched with AT→GC SNVs than in Native Americans. Furthermore, we showed that deleterious homozygous SNVs are also predominantly AT→GC in Africans, and to a greater extent than in non-Africans. This suggests the possibility that recessive genetic disorders in Africans are more likely to be caused by AT→GC variants than in non-Africans. Therefore, our study recommends that genome-wide association studies should consider the frequency of population specific nucleotide changes.

## Materials and Methods

### Genome data

We obtained genotype data from the 1000 genome project – Phase II (Genomes Project, et al. 2015). The genome-wide variations from the 26 populations including Africans (seven populations: 661 individuals), South Asians (five: 489), European (five: 503), East Asian (five: 504) and South Americans (four: 347). Although there were 85 Peruvian genomes were available, most of these were admixed with Europeans and Africans. Hence, we used the likelihood based clustering algorithm *Admixture* (Alexander, et al. 2009) and examined the proportion of admixture in each Peruvian genome. Our results showed that only 20 genomes (40 chromosomes) had <0.5% admixture from other populations and we included these un-admixed Peruvians as the 27^th^ population in our analyses. To identify derived alleles, orientations of SNVs were determined using the ancestral state of the nucleotides, which was inferred from six primate EPO alignments (Genomes Project, et al. 2010). The SNVs were divided into eleven categories based on their derived allele frequencies (see Figure 2-legend). For the SNVs in each category we computed the counts of six types of changes: A/T→G/C, G/C→A/T, C→G, G→C, A→T and T→A (see below).

We also obtained the genotype data from the Simon Genome Diversity Project (Mallick, et al. 2016). To examine the patterns of nucleotide changes we used the homozygous and heterozygous SNVs present in a single representative genome from each of the 126 populations. We excluded four African hunter-gatherer populations from our analysis as it was difficult to ascertain the correct orientation of the nucleotide changes in these genomes. For each genome, we estimated the number of homozygous and heterozygous changes belonging to the six types described above.

### Deleterious mutations

To determine the deleterious nature of a SNV we used a robust method, Combined Annotation-Dependent Depletion (CADD) that integrates diverse annotations into a single measure (C score) (Kircher, et al. 2014). The extent of deleteriousness was further determined by estimating the corresponding selective coefficients for these scores (Racimo and Schraiber 2014). The C scores for each SNV in the 1000 Genomes Project data were publically available (http://cadd.gs.washington.edu/download). Using an in-house Perl script, we combined this score with the genome data by using the chromosomal co-ordinates of the SNVs. For the deleterious variant analysis, we included only the SNVs for which the C score was available. We used a C score of ≥15 to determine a mutation to be deleterious in nature following previous studies (Kircher, et al. 2014; Subramanian 2015). However, using a different threshold produced almost identical results.

### Estimating the ratio of mutation rates

The ratio of AT→GC and GC→AT mutation rates could be estimated based on the Waterson estimator (Watterson 1875), *θ* = 4*N*_*e*_*μ* = *S*/*a*_*n*_, where *N*_*e*_ is the effective population size and *S* is the number of segregating sites per site and 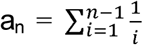. We can use this estimator considering only one type of mutation as:

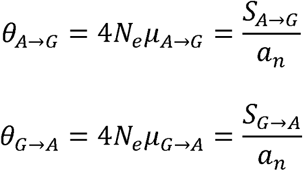

The ratio of forward and reverse nucleotide changes (*β*) could be obtained as:

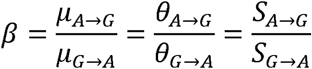

The number of segregating sites per site or SNVs per site of a genome can be estimated as:

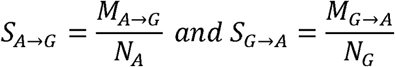

where *M*_A→G_ and *M*_G→A_ are the number of observed A→G and G→A mutations in a genome respectively and *N*_A_ and *N*_G_ are the number of ancestral A and G nucleotides. This formula can be extended for the combined AT→GC and GC→AT mutation rates because each pattern is mutually exclusive and hence the ratio of nucleotide changes (*β*) is:

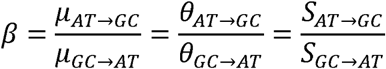

The number of segregating sites or SNVs in a genome can be calculated as:

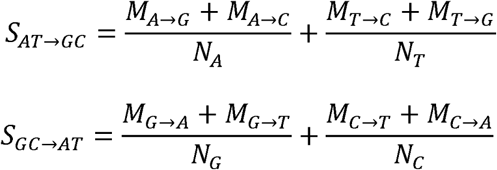

Since A and T as well as C and G are complementary to each other in a double-stranded DNA they are equal in number. Therefore, *β* can be expressed as,

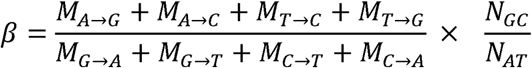

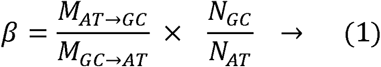

This derivation proves that the ratio of forward and reverse rates of changes can be calculated by simply taking the ratio of the observed counts of AT→GC (*M*_*AT→GC*_) and GC→AT (*M*_*GC→AT*_) changes and multiplying with the ratio of the number of GC (*N*_GC_) to AT (*N*_AT_) nucleotides in a genome. Since equation 1 represents the ratio of mutation rates (*μ*_*AT→GC*_ and *μ*_*GC→AT*_) this ratio is expected to be 1 (*β* =1) if the observed nucleotide changes are solely due to the result of these mutation rates. Any deviation from this suggests a bias in the substitution process. While *β* >1 indicate an excess of AT→GC substitutions *β* <1 imply an excess of GC→AT substitutions.

GC-biased gene conversion is known to affect only the changes involving weak (A or T) to strong (G or C) nucleotides but not the changes within weak (A↔T) or within strong (G↔C) nucleotides. Hence the latter is not expected to vary between populations with different *N*_*e*_. Therefore, we used this as a normalization factor and developed a normalized ratio of AT→GC to GC→AT (*β’*).

The A→G mutation rate can be normalized using A→T rate and the normalized rate (*τ*_A→G_) can be expressed as:

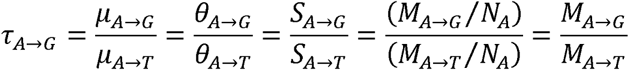

Similarly, we can obtain this expression for AT→GC and GC→AT rates as:

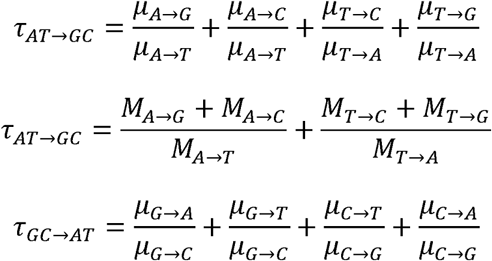

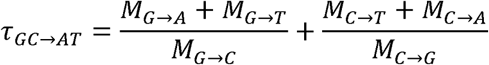

Therefore, the normalized ratio of AT→GC to GC→AT (*β’*) is:

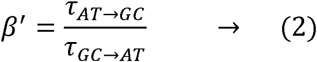

The relationship between nucleotide diversity (*π*), mutation rate (*μ*) and effective population size (*N*_*e*_) for diploid organisms is *π* = 4*N*_*e*_*μ*. Using this relationship, we calculated the effective population size as *N*_*e*_ = *π*/4*μ*. We used the observed nucleotide diversity of a population or of a diploid genome and used a mutation rate of 1.2 × 10^-8^ substitutions per site per generation following many studies based on human pedigree genome data (Roach, et al. 2010; Conrad, et al. 2011). A recent suggested that the rate of mutation accumulation in non-European genomes could be slightly (5%) higher than that of Africans (Mallick, et al. 2016). To accommodate this difference, we subtracted 5% of the nucleotide diversity for non-African populations only while calculating the effective population sizes and obtained almost identical results (Supplementary figure S4).

### Estimation of the proportion of AT→GC counts

We also estimated the proportion of AT→GC counts (*P*_*GC*_) and GC→AT (*N*_*GC*→*AT*_) counts, which were calculated as:

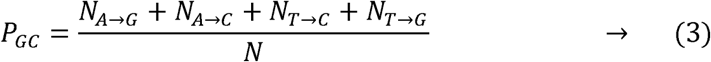

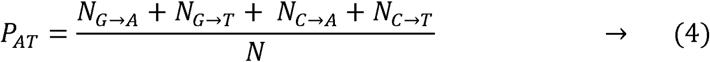

*N* is the number of all types of nucleotide changes. The standard error of *P*_*AT*→*GC*_ and *P*_*GC*→*AT*_ were calculated using the binomial variance.

## Acknowledgment

The author thanks Alex Quin for critical comments and acknowledges the support from the Australian Research Council (LP160100594) and the University of the Sunshine Coast.

## Author Contributions

Conceptualization: SS

Data Analysis: SS

Writing: SS

